# Distributed encoding of hippocampal information in mossy cells

**DOI:** 10.1101/2024.03.14.584957

**Authors:** Ayako Ouchi, Taro Toyoizumi, Yuji Ikegaya

**Affiliations:** Graduate School of Pharmaceutical Sciences, The University of Tokyo, Tokyo 113-0033, Japan; Laboratory for Neural Computation and Adaptation, RIKEN CBS, Wako, Saitama, 351-0198, Japan; Institute of AI and Beyond, The University of Tokyo, Tokyo 113-0033, Japan; Center for Information and Neural Networks, National Institute of Information and Communications Technology, Suita City, Osaka, 565-0871, Japan

**Author notes:** To whom correspondence should be addressed: Yuji Ikegaya, Ph.D., Laboratory of Chemical Pharmacology, Graduate School of Pharmaceutical Sciences, The University of Tokyo 7-3-1 Hongo, Bunkyo-ku, Tokyo 113-0033, Japan.

**Keywords:** hippocampus, sharp wave, ripple, mossy cell, dentate gyrus

## Abstract

In neural information processing, the nervous system transmits neuronal activity across layers of neural circuits, occasionally passing through small layers composed only of sparse neurons. Hippocampal hilar mossy cells (MCs) constitute such a typical bottleneck layer. *In vivo/vitro* patch-clamp recordings revealed that MCs were reliably depolarized in response to sharp-wave ripples (SWRs), synchronous neuronal events transmitted from the CA3 region to the dentate gyrus via the MC layer. Machine-learning algorithms predicted the waveforms of SWRs in the CA3 region, based on the MC depolarization waveforms, suggesting that CA3 neural information is indeed transmitted to the MC layer. However, the prediction accuracy varied; *i.e.*, a particular MC showed a more robust association with a particular SWR cluster, and the SWR cluster associated with one MC rarely overlapped with the SWR clusters associated with other MCs. Thus, CA3 network activity is distributed across MC ensembles with pseudo-orthogonal neural representations, allowing the small MC layer to effectively compress hippocampal information.

## Introduction

Neural information is processed as it travels through the various layers of the neural circuit, with the number of neurons in each layer varying from layer to layer. In general, a layer with a large number of neurons is assumed to be able to process larger and more complex information, while a layer with a small number of neurons must compress or otherwise trim the information. Neural processing with a small layer may be advantageous for dimensionality and feature extraction of information (Grezl et al., 2007).

Mossy cells (MCs) in the hippocampal formation serve as a good experimental model for such a bottleneck middle layer (**Figure S1**). They are the only excitatory neurons that exist in the dentate hilus, a small zone intercalated between the CA3 region and the dentate gyrus (DG). MCs receive synaptic inputs from CA3 pyramidal cells, and their axons send synaptic outputs to DG granule cells. Thus, MCs relay neuronal activity from the CA3 region to the DG; note that there is no direct axonal projection from CA3 pyramidal cells to DG granule cells (Fujise and Kosaka, 1999). An estimated 15,000 MCs reside in the rat hippocampal formation, whereas approximately 300,000 and 2,450,000 CA3 pyramidal and granule cells exist, respectively (Amaral et al., 1990; Jinno and Kosaka, 2010). Thus, the number of MCs is one to two orders of magnitude lower than that of CA3 pyramidal neurons and DG granule cells (**Figure S1**), and MCs constitute a small relay layer.

Consistent with these anatomical features, sharp wave-ripples (SWs) propagate from the CA3 region to the DG via MCs (Penttonen et al., 1997; Scharfman, 2007). SWs are high-frequency oscillations (80-200Hz) in the local field potential (LFP) of the hippocampus that occur during sleep or quiet wakefulness (Buzsáki et al., 1992). SWs represent a mechanism for reactivating or consolidating recent experiences, allowing the ‘replayed’ information to be transferred from the CA3 region forward to the neocortex via the subiculum (Nitzan et al., 2020) as well as backward to the DG. The CA3-DG backpropagation of SWs is experimentally captured by the depolarization of MCs during SWs (Ouchi et al., 2017; Swaminathan et al., 2018) and is theoretically thought to play an important role in pattern separation (Myers and Scharfman, 2011).

During a SW, spatiotemporal patterns of neuronal firing in the hippocampus that resembles activity during an experience is replayed as rapid spike sequences (Diba and Buzsáki, 2007; Lee and Wilson, 2002). Different SWs may reactivate different sequences of spikes, possibly encoding information that correlates with different experiences. Neural information in SWs is manifested in the LFP waveforms of SWs (Navas-Olive et al., 2022; Taxidis et al., 2015). Accordingly, the characteristics of SW waveforms, including their amplitudes and widths, exhibit heterogeneity across individual SWs. Therefore, comparing the SW waveforms with SW-induced depolarizations in MCs provides us with a unique opportunity to investigate how neural information travels through a small layer.

In the present work, we performed patch-clamp recording from up to 5 MCs together with recording of CA3 SWs and developed a neural network that associates SWs with membrane potential (*V*m) dynamics in MCs. Our neural network was able to predict SW waveforms from MC *V*ms. Strikingly, about 9% of the total SW waveform could be predicted from even a single MC. Neural representations of MCs cover a wide range of the SW repertoire with slight overlap between MCs, and as a result, the proportion of predictable SWs increased sublinearly as the number of MCs used for prediction increased.

## Results

### Simultaneous patch-clamp recordings from hilar MCs *in vitro* and *in vivo*

Using mouse acute hippocampal slice preparations, we simultaneously recorded *V*m from multiple neurons in the hilus in a whole-cell current-clamp configuration as well as local field potentials (LFPs) from the CA3 pyramidal cell layer (**Figure 1A**). A post hoc biocytin-based visualization method was used to confocally image the recorded neurons in NeuroTrace Nissl-counterstained slices (**Figure 1A**, inset). Data were excluded from analysis if the recorded cells had a membrane capacitance lower than 45 pF (Hedrick et al., 2017) and had spines and thorny excrescences on their proximal dendrites (Murakawa and Kosaka, 2001), which are electrophysiological and morphological markers of MCs. As a result, we obtained 2 quintuple, 8 quadruple, and 15 triple patch-clamp recordings from MCs in 23 slices from 14 mice. The recording periods ranged from 55 s to 357 s (median = 301 s).

**Figure 1.**
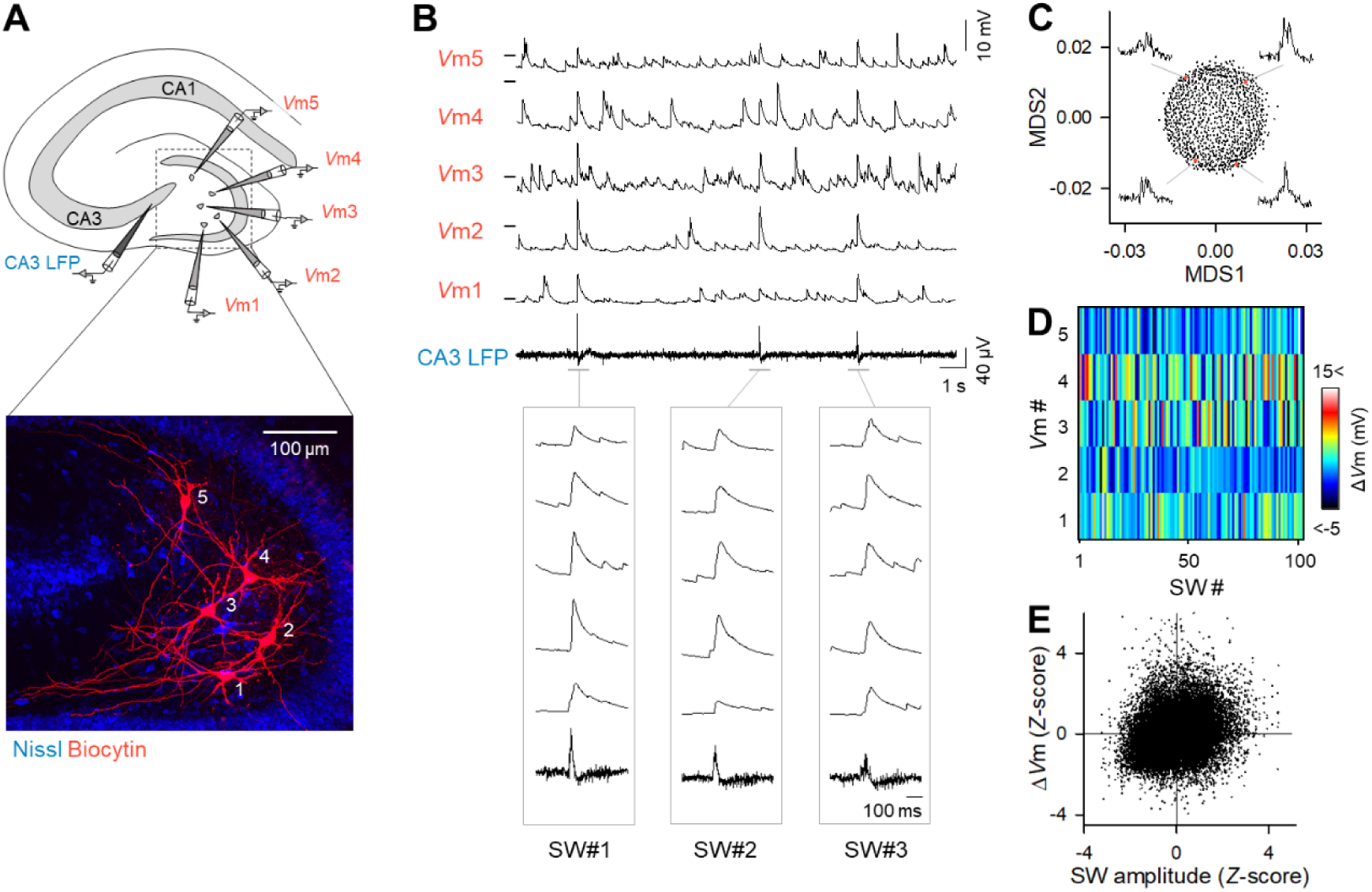
Spatiotemporal diversity of CA3 SW-induced Δ*V*ms in MCs. (**A**) (Top) Schematic illustrations of whole-cell *V*m recordings from five MCs together with LFP recordings from the CA3 region. (Bottom) Recorded cells were confocally identified by immunostaining for intracellularly injected biocytin (red) in a Nissl-counterstained slice (blue). (**B**) Representative traces of five MC *V*ms and CA3 LFPs. The horizontal ticks on the left of the Vm traces represent −60 mV. The traces during the three SWs are expanded in time in the bottom boxes. (**C**) The LFP traces of 945 SWs in a representative dataset were dimensionally reduced using the MDS algorithm. Each dot indicates a single SW event. (**D**) A pseudocolor map for the amplitudes of the SW-induced depolarizations (Δ*V*m) in five MCs during a total of 102 SWs, demonstrating the heterogeneity of *V*m responses among MCs across SWs. (**E**) Weak correlations between the SW amplitudes and Δ*V*ms of MCs. Each dot indicates a single SW event. Δ*V*ms and SW amplitudes were *Z*-standardized for each recording electrode before the data were pooled. *R* = 0.25, *n* = 28,463 SW events in 87 mossy cells from 23 slices.

All tested slices spontaneously emitted SWs in CA3 LFPs (**Figure 1B**). The frequency of SW events ranged from 11 to 84 per minute (median = 38 per minute). The waveforms of SWs were not uniform; for example, their amplitudes and widths varied from SW to SW. Dimension reduction using the multidimensional scaling (MDS) algorithm revealed that the SW waveforms formed a continuous set, because they were nondiscretely distributed in two-dimensional MDS space (**Figure 1C**).

MCs reliably exhibited depolarization immediately after SW onset (**Figure 1B**). This reliability is in marked contrast to that of the CA1 pyramidal cells and DG granule cells, which do not always depolarize for all SWs (Mizunuma et al., 2014; Ouchi et al., 2017). The *V*m responses of MCs were abolished when the dentate hilus was surgically isolated from the CA3 region (**Figure S2**), indicating that these responses were caused by CA3 SW activity.

The *V*m responses of MCs to SWs were also heterogeneous; a single MC exhibited different depolarization amplitudes (Δ*V*ms) for different SW events, whereas a single SW induced different Δ*V*ms in different MCs (**Figure 1D**). Overall, the Δ*V*ms in MCs were only weakly correlated with the amplitude of SWs (**Figure 1E**; *R* = 0.25, *n* = 28,463 SWs from 87 cells), indicating that even a small SW could cause a large Δ*V*m in a MC and vice versa.

We also performed *in vivo* whole-cell recordings from MCs in urethane-anesthetized mice (**Figure 2A**). The recorded cells were identified based on their morphological features and immunoreactivity for glutamate receptor 2/3 (GluR2/3), a molecular marker of MCs (**Figure 2B**). As observed *in vitro*, MCs exhibited diverse Δ*V*m responses to SWs (**Figure 2C**), and again, Δ*V*ms correlated only weakly with SW amplitudes (**Figure 2D**; *R* = 0.1053, *n* = 756 SWs from 6 cells).

**Figure 2.**
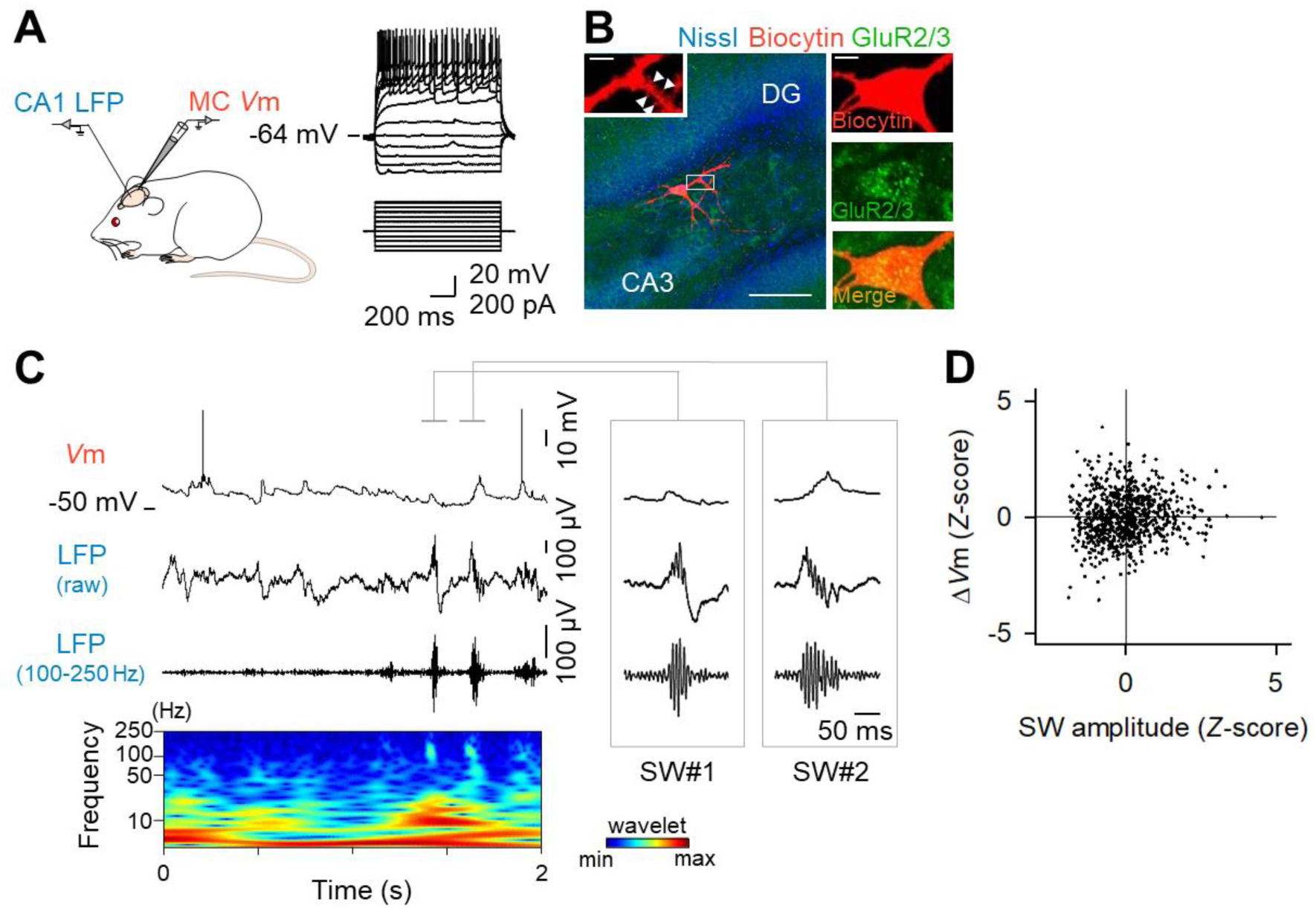
Diversity in SW-induced MC Δ*V*ms *in vivo.* (**A**) MCs were current-clamped in a urethane-anesthetized mouse, while the LFPs were recorded from the CA1 region. (**B**) The recorded neurons were confocally visualized by the detection of intracellularly loaded biocytin (red) in a section immunostained with anti-GluR2/3 (green), a MC marker, and counterstained with NeuroTrace Nissl (blue). Scale bar: 100 µm. The high-magnification images on the right indicate that the recorded neuron was positive for GluR2/3 and that thorny excrescences were observed on proximal dendrites. Scale bar: 10 µm. (**C**) (Top) Representative *V*m traces of a MC. (Middle) A raw LFP trace recorded simultaneously from the CA1 region and its bandpass-filtered trace (100-250 Hz). The bottom panel indicates the wavelet spectrogram of the raw LFP trace. The traces during two SWs are expanded in time in the right insets. (**D**) Weak correlations between the SW amplitudes and Δ*V*ms of MCs during SWs. *R* = 0.1053, *n* = 756 SW events from 6 MCs.

The weak correlations between Δ*V*ms and SW amplitudes suggest that MCs respond to SWs in a complex, rather than a simple linear, manner. The variability in Δ*V*ms responses suggests that MC ensembles may collectively cover information of diverse SWs. Therefore, we hypothesized that, if this is the case, SW waveforms (SW-related information) could be predicted from the *V*m responses of a set of MCs.

### Machine learning-based prediction of SW traces from *V*ms in MCs

To associate MC *V*ms with SW waveforms, we trained a neural network model with a hidden layer (**Figure 3A**). For each SW event in quintuple patch-clamp recordings, we extracted 100-ms segments of the *V*m and LFP traces between −50 and +50 ms relative to the SW peak times. We divided these segments into 5 subsets and trained the neural network using *V*m and LFP traces in the 4 out of 5 subsets (80%) to predict the SW waveforms in the remaining subset (20%) that was not used for training (**Figure 3B**). We repeated this procedure so that all SWs were targeted for prediction. For each prediction, the prediction error was quantified by the root mean square error (RMSE) between the original and predicted SW waveforms. We also trained the same neural network using surrogate data in which the combinations of *V*ms and LFPs were randomly shuffled across SW events within each recording dataset. We then compared the RMSEs for the LFP waveforms predicted by the original and shuffled *V*m data. The RMSEs from a total of 271 SWs from 2 quintuple datasets were pooled for a cumulative plot (**Figure 3C**). The RMSEs produced by prediction using the real data were significantly lower than those produced by the shuffled data (*P* = 0.0026), indicating that the neural network predicted the SW waveforms from the *V*m dynamics of five MCs with significantly higher accuracy than chance. In other words, CA3 information during SWs was at least partially preserved in the *V*m responses of MCs.

**Figure 3.**
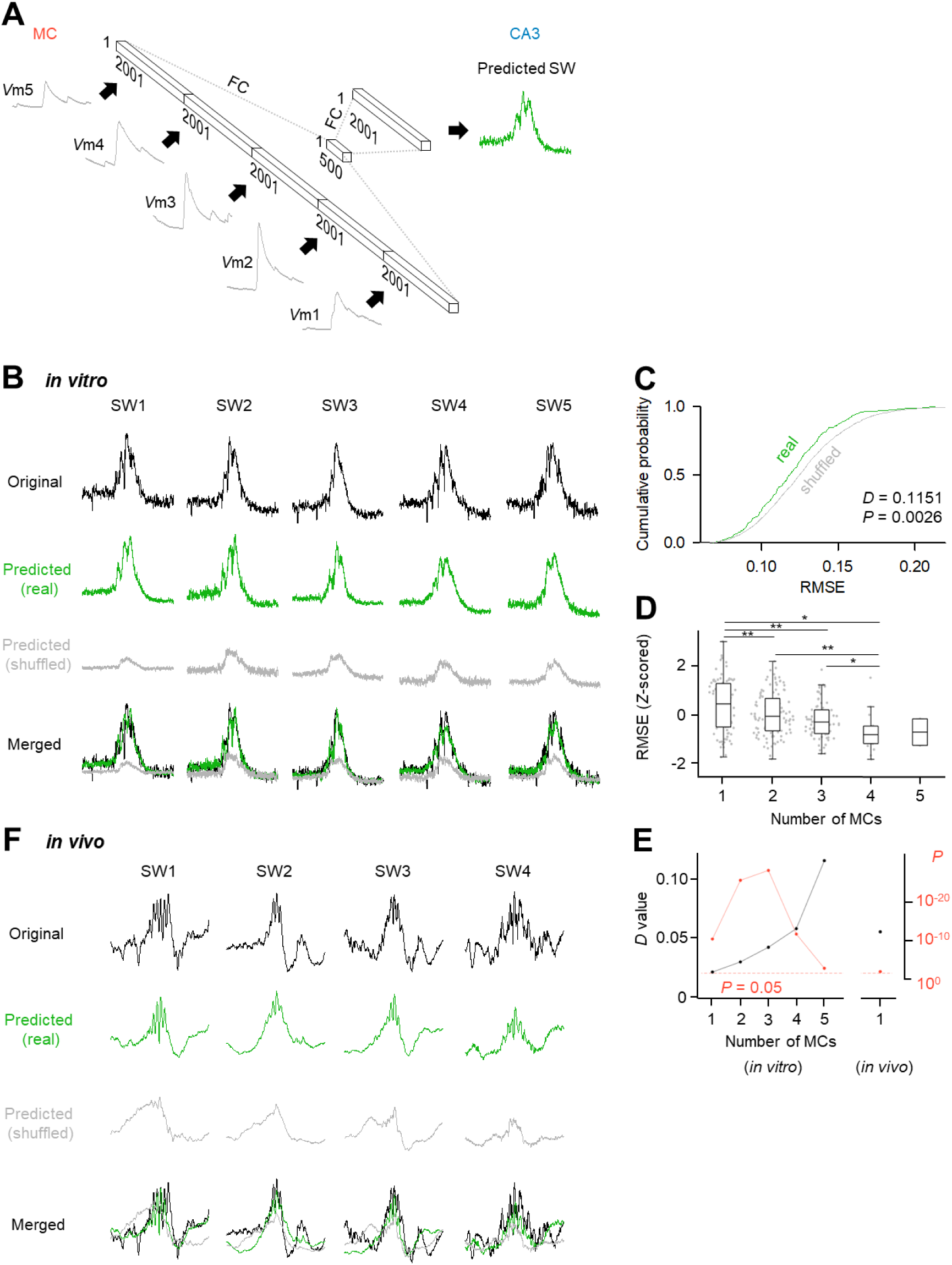
Machine learning-based prediction of SW traces from *V*ms in MCs *in vitro*/ *in vivo*. (**A**) Design of the neural network, which consisted of five layers connected unidirectionally with fully connected (FC) links. (**B**) Five examples of original SW traces (black) and SW traces predicted from real *V*m datasets (green) and shuffled *V*m datasets (gray). All traces are merged at the bottom. (**C**)The cumulative distribution of the RMSEs between the original and predicted traces (prediction errors) indicates that real dataset-based prediction had significantly smaller prediction errors than the shuffled dataset-based prediction. *n* = 271 SWs from 2 recording datasets of MC quintets, \**P* = 0.0028, *D* = 0.1151, Kolmogorov–Smirnov test. (**D**) RMSE calculated between the original and predicted traces were decreased as the number of simultaneously recorded cells used for prediction increased. One cell: *n* = 28,463 SWs from 87 single MC recordings; 2 cells: *n* = 35,914 SWs from 113 MC pair recordings; 3 cells: *n* = 19,874 SWs from 67 trios; 4 cells: *n* = 4,403 SWs from 18 quartets; 5 cells; *n* = 271 SWs from 2 quintets, *F* = 8.28, *P* = 2.51 × 10^-6^, one-way ANOVA. \**P* < 0.05, \*\**P* < 0.005, Fisher’s LSD test. (**E**) The *D* values determined by the Kolmogorov–Smirnov test (black) increased as the number of simultaneously recorded cells used for prediction increased. The *P* values for the corresponding *D* values are plotted in *red*, indicating that the *V*ms of 1-5 MCs led to a significantly accurate prediction. The *D* value and *P* value were also calculated from *in vivo* dataset as in Fig. 3C and plotted on the right. (*in vitro*) One cell: *n* = 28,463 SWs from 87 single MC recordings; 2 cells: *n* = 35,914 SWs from 113 MC pair recordings; 3 cells: *n* = 19,874 SWs from 67 trios; 4 cells: *n* = 4,403 SWs from 18 quartets; 5 cells; (*in vivo*) *n* = 273 SWs from 2 quintets. *n* = 765 SWs from 6 mice, \**P* = 0.0185, *D* = 0.0553, Kolmogorov–Smirnov test. (**F**) Four examples of original SW traces (black) and SW traces predicted from real *V*m datasets (green) and shuffled *V*m datasets (gray). All traces are merged in the bottom panel.

We next investigated the relationship between the number of MCs used to train the neural network and the prediction performance. To increase the number of datasets, we divided the quintuple, quadruple, and triple patch-clamp recording datasets into subsets with fewer MCs. The RMSEs calculated between the original and predicted traces, which were *Z*-standardized for each dataset, decreased as the number of simultaneously recorded cells used for prediction increased (**Figure 3D**). The prediction performance was also evaluated using the *D* value of the Kolmogorov-Smirnov test. The *D* values increased as the number of simultaneously recorded cells increased (**Figure 3E**). As another index of prediction accuracy, we calculated the wavelet coherence for a high-frequency range (120-250 Hz) between the original and predicted waveforms and found that the high-frequency details of the SWR waveforms predicted by a larger number of MCs were also closer to the original (**Figure S3**), suggesting that a greater number of MCs can retain more information about SWRs. Remarkably, the *V*ms of even a single MC were sufficient to significantly predict the SW waveforms (**Figure 3E**; *D =* 0.0215, *P* = 7.09×10^-11^, *n* = 87 cells). We also trained the neural network using *V*m data obtained by *in vivo* single-cell patch-clamp recordings to predict the LFP waveforms during SWs (**Figure S4**; **Figure 3F**). Again, the RMSEs predicted by the real datasets were significantly smaller than those predicted by the shuffled datasets (**Figure 3E**, *D =* 0.0553, *P* = 0.0185, *n* = 6 cells).

### Proportion of predictable SWs by single MC and correlation with spatial distribution or fundamental neuronal properties of MCs

Given that the SW waveform was predicted with statistical significance even with a single MC, we examined the percentage of SWs that a single MC could predict (predictable SWs) relative to the total SWs. For each MC, the significance of the RMSEs in predicting individual SW waveforms was statistically determined (**Figure 4A**); Specifically, if the RMSE predicted by the real data sets (RMSE_real_) was significantly lower than that predicted by the shuffled data sets (RMSE_surrogate_), where RMSE_real_ and RMSE_surrogate_ represent the RMSEs between the original SW waveform and the SW waveforms predicted by the neural networks trained with the real *V*ms of the MC and its surrogate *V*ms, respectively, then the SW was defined as ‘predictable’ by the MC. The surrogate *V*ms were generated by randomly shuffling *V*ms across SWs within the MC. The percentage of predictable SWs to all SWs recorded from the MC was 9.2 ± 4.8% (mean ± SD of 87 MCs, median = 8.01). The relative positions of individual MCs in the dentate hilus were plotted on a standardized semicircular map, with the predictable SW ratios of the corresponding MCs displayed in pseudocolor (Fig. 4B) (Koyama et al., 2012). We computed the spatial entropy of this distribution to examine whether MCs with specific SW predictability were spatially clustered in a particular area. The spatial entropy was 6.39, and this value was significantly lower than the 95% confidence interval of a chance distribution computed from 10,000 proxy data sets in which the percentage of predictable SWs was randomly shuffled across MCs without changing the positions of the MCs (**Figure 4C**, *P* = 4.0×10^-4^). Therefore, the anatomical locations of MCs were linked with their percentage of the predictable SWs; specifically, the lower blade of the DG included MC with higher predictability. However, neither the membrane capacitances (**Figure 4D**) nor the membrane resistances (**Figure 4E**) of MCs were correlated with the percentage of predictable SWs of the MCs. Thus, the intrinsic electrophysiological properties of MCs were unlikely to determine their SW predictability.

**Figure 4.**
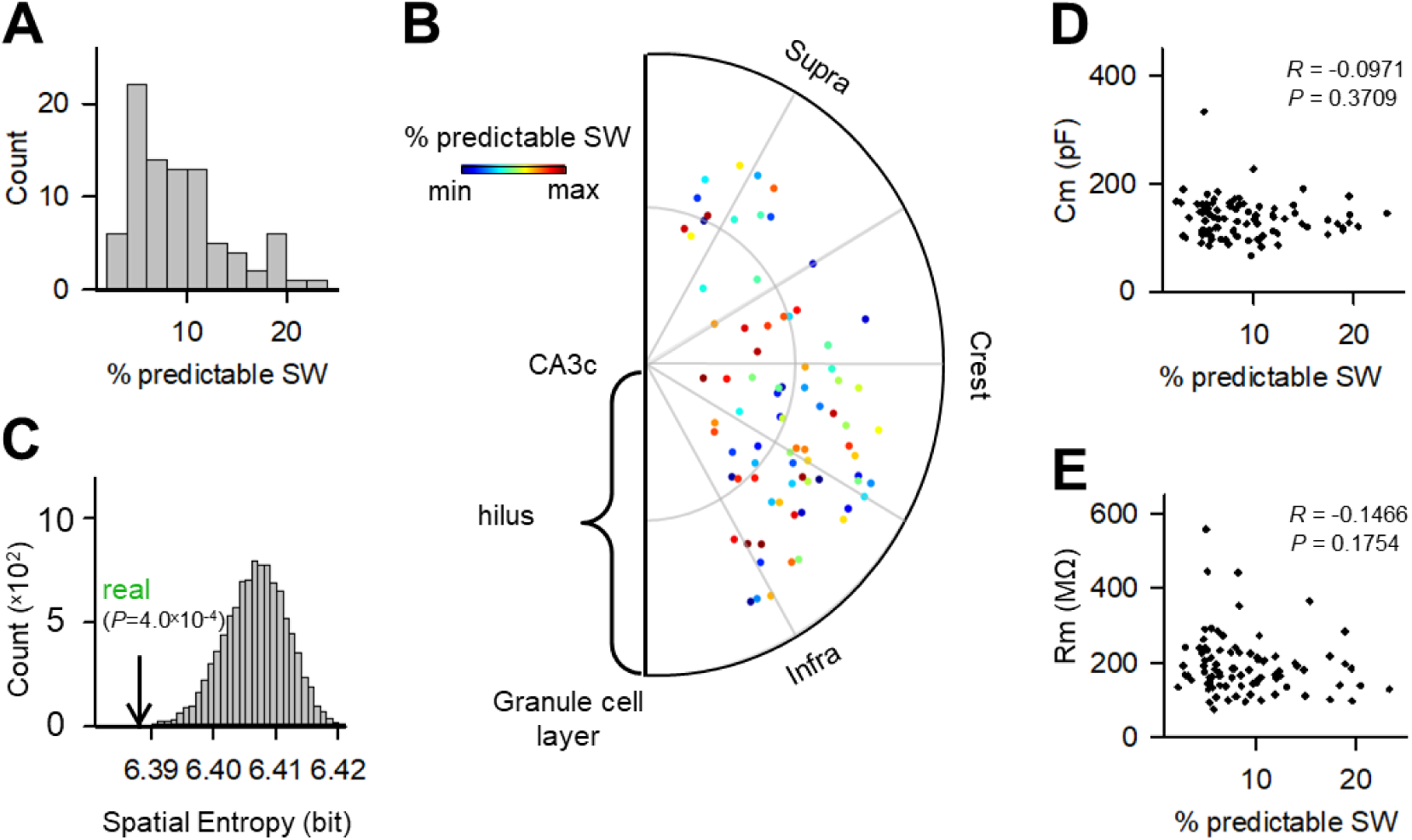
Correlation between proportions of predictable SW by single MC and spatial distribution or fundamental neuronal properties. (**A**) Distribution of the % predictable SW from 87 MCs. The % predictable SW was computed from RMSEs by using a single MC prediction. (**B**) The relative locations of MCs in the hilus are plotted in a semicircular diagram. Each dot indicates a MC, and its color represents the % predictable SW. (**C**) Spatial bias of (B) was evaluated by spatial entropy and compared to its chance distribution. The real entropy value is indicated by an arrow. (**D, E**) Correlations between % predictable SW and electrophysiological properties. Each dot indicates a single SW. Cm: *P* = 0.3709, *R* = −0.0971; Rm: *P* = 0.1754, *R* = −0.1466, *n* = 87 MCs.

### Prediction deflection of SWs in MCs

We investigated whether SWs with a given waveform were more predictable by a given MC. The dimensionality of the SW waveforms was reduced to two dimensions using the MDS algorithm. Because the MDS algorithm preserves the relative distances (here, RMSE values) between any two SWs, the proximity of SW pairs in MDS space reflects the similarity of their original waveforms. As an example, we plotted a total of 945 SWs recorded from a hippocampal slice in which four MCs were simultaneously patch-clamped (**Figure 5A**, left). The predictable SWs are shown in red in the MDS space in **Figure 5A**. We then computed the spatial entropy of the predictable SWs in the MDS plot and compared it to the chance distribution obtained by 1,000,000 surrogates in which the binarized prediction (*i.e.*, predictable or non-predictable) was permuted across SWs within the cell (**Figure 5A**, right). We repeated this comparison for a total of 87 MCs and found that for 37 out of 87 MCs, the spatial entropy was significantly lower than the chance value (**Figure 5B**). Thus, as a whole, a MC tended to predict a specific subset of SWs with similar waveforms.

**Figure 5.**
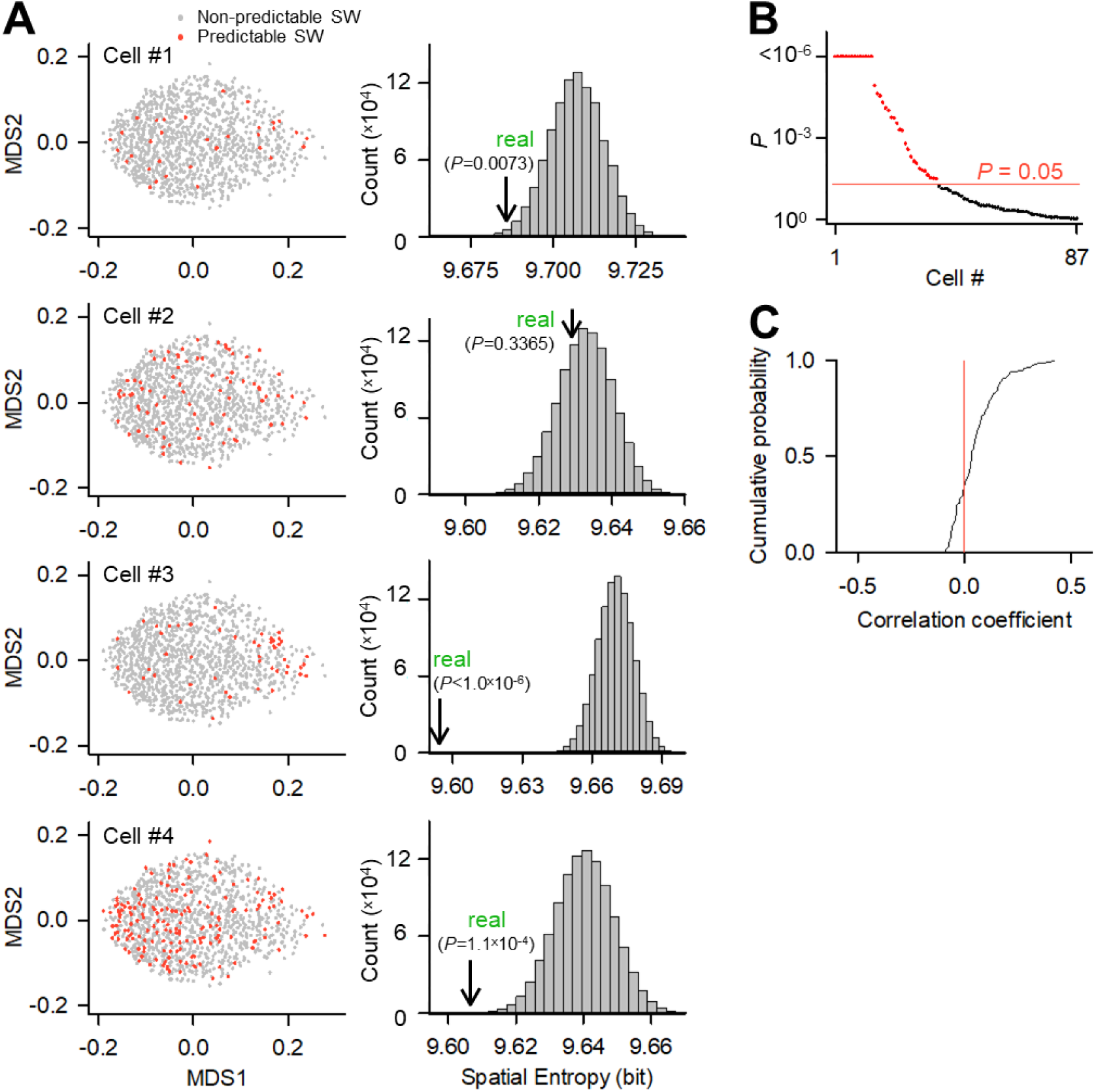
Predicted deflection of SWs in MCs. (**A**) (Left) The LFP traces of 945 SWs from a representative quadruple recording dataset were dimensionally reduced using the MDS algorithm. Each dot indicates a single SW event, and its color represents whether the prediction rate is significant (red) or not (gray). (Right) The spatial bias of the prediction rates in the MDS space was evaluated by spatial entropy. Lower entropies indicate higher spatial biases of the prediction rate. In each MC, the entropy was compared to the change distribution obtained by 1,000,000 surrogates in which the prediction rates were shuffled within the MC. The real entropy value is indicated by an arrow with its *P* value. (**B**) The same calculations as those in (A) were repeated for all 87 cells, and their *P* values were plotted. The red dots indicate MCs with significantly lower entropy values. (**C**) Distribution of the correlation coefficient of the RMSE score sets (Non-predictable SW as 0, Predictable SW as 1) between 113 MC pairs, which were obtained from 2 quintuple, 8 quadruple, and 15 triple recording datasets.

We examined how different MCs in a hippocampal slice predicted the different SW subsets. For this purpose, we considered the binarized prediction scores for SWs as a vector. Specifically, with ‘predictable’ as 1 and ‘non-predictable’ as 0, we created a binary vector for all SWs in a given MC. We then calculated the correlation coefficient between the vectors of two MCs. Data were pooled from a total of 113 MC pairs from 2 quintuple, 8 quadruple and 15 triple recording datasets. The correlation coefficients varied mainly between 0 and 0.3, and their mean (0.054) was relatively small with SD = 0.105. Indeed, they did not differ from 0 as a whole distribution (*Z =* −0.002, *P* = 0.998, *Z* test for single value comparison), indicating that a subset of SWs that can be predicted by a given MC is nearly independent of that predicted by the other MCs (**Figure 5C**). Therefore, information in CA3 SWs is represented in a manner that is nearly mutually exclusive (*i.e.*, pseudo-orthogonal) among individual MCs.

### Prediction specificity of SWs by MCs

Further analysis was performed to confirm the pseudo-orthogonality of the information code. Binary vectors were computed in the analysis of **Figure 5A**. The vectors for five representative MCs recorded from the same hippocampal slice containing a total of 178 SWs are shown in **Figure 6A**, indicating that sometimes different MCs were able to predict the same wave, sometimes not. The Venn diagram shows how the SWs predicted by these 5 MCs overlapped (**Figure 6B**). The number of SWs in each area of the Venn diagram is shown in Fig. S5. These data suggest that the more MCs recorded simultaneously, the greater the total number of predictable SWs. Indeed, the pooled plot of a total of 88,925 SWs recorded from 23 slices shows that the percentage of predictable SWs increased as the number of simultaneously recorded MCs increased (**Figure 6C**; *P* = 1.00 × 10^-38^, Jonckheere-Terpstra test). However, predictable SW did not increase completely linearly with the number of MCs, but tended to increase sublinearly. This sublinearity arises because there exist a small number of cases where multiple MCs predicted the same SW (overlapping SW). We calculated the percentage of overlapping SWs that were predictable by both of any two MCs to the total SWs for each recording data set (**Figure 6D**; median = 1.56, SD = 2.04, *n* = 113 MC pairs). The percentage of overlapping SWs was sparse from record to record but never exceeded 10%, and this result confirms again that overlapping SWs were in the minority.

**Figure 6.**
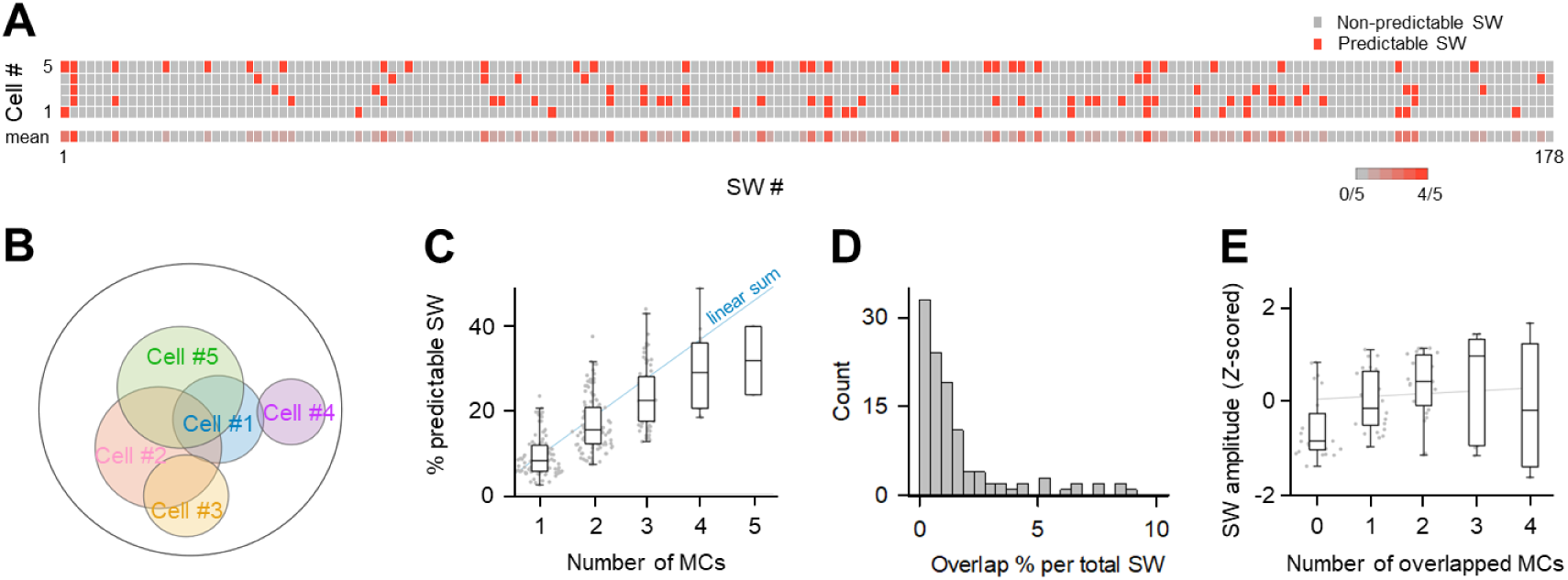
Prediction specificity of SWs by MCs. (**A**) The representative quintuple recording in a total of 178 SWs. The top five rows demonstrating the SWs that each MC can predict. Its color represents whether the SW was predictable (red) or not-predictable (gray). The mean predictivity across five MCs is shown in the bottom rastergram. (**B**) The Venn diagram demonstrated the image of the range of SWs that each MC can predict out of the recorded SW. (**C**) The % predictable SW was increased as the number of simultaneously recorded cells used for prediction increased. The linear summation is indicated by solid line (blue). One cell: *n* = 28,463 SWs from 87 single MC recordings; 2 cells: *n* = 35,914 SWs from 113 MC pair recordings; 3 cells: *n* = 19,874 SWs from 67 trios; 4 cells: *n* = 4,403 SWs from 18 quartets; 5 cells; *n* = 271 SWs from 2 quintets, *\*P* = 1.00 × 10^-38^, Jonckheere-Terpstra test. (**D**) Distribution of the overlap % per total SW between 113 MC pairs, which were obtained from 2 quintuple, 8 quadruple, and 15 triple recording datasets. (**E**) Correlations between SW amplitudes and number of overlapped MCs. The regression line is indicated by solid line (gray). The data were obtained from 2 quintuple, 8 quadruple, and 15 triple recording datasets, *F* = 5.38, \**P* = 0.0007, one-way ANOVA; *\*P* = 1.59 × 10^-5^, Jonckheere-Terpstra test.

To characterize the overlapping SWs, we analyzed their waveforms. The amplitudes of overlapping SWs had a slight but significant positive correlation with the number of MCs that could predict the SWs (**Figure 6E**, *P* = 1.59×10^-5^, Jonckheere-Terpstra test), suggesting that smaller SWs are covered by a smaller number of MCs, while larger amplitude SWs are covered by a larger number of MCs. Other waveform properties, including the duration of SWs, the peak oscillation frequency of ripples during SWs, the Fast Fourier Transform (FFT) power, and the area under the curve (AUC) of SWs, did not depend on the number of predicting MCs. (**Figure S6**).

## Discussion

In the present study, we performed simultaneous *in vitro* whole-cell recordings of up to five hilar MCs in hippocampal slices and *in vivo* whole-cell recordings of single MCs from anesthetized mice and found that MCs responded to SWs with their *V*ms, which could predict the waveforms of SWs in the CA3 region. The accuracy of predicting SW waveforms improved with the number of simultaneously recorded MCs, but the *V*ms of even a single MC could predict SWs more accurately than expected by chance. Predictability was heterogeneous among SWs, but a subset of SWs predicted by one MC differed greatly from those predicted by another MC, and only a small fraction of SWs were predictable by multiple MCs. As a result, the proportion of predictable SWs increased efficiently with the number of simultaneously recorded MCs, and more than 30% of the total SWs were predictable when five MCs were recorded.

In this study, we used a novel mathematical approach to correlate the upstream neural dynamics with the downstream neural response. In general, it is difficult to elucidate the relationships between the intracellular activity of neurons and the extracellular activity of neurons. In our preliminary studies, we performed several traditional experiments aimed at identifying a function that could correlate multiple *V*ms with LFPs. However, conventional multivariate and regression analyses proved insufficient to describe LFPs in relation to *V*ms. We therefore designed a neural network that was inspired by the autoencoder networks (Hinton and Salakhutdinov, 2006). The autoencoders have a bottleneck hidden layer that extracts low-dimensional codes in high-dimensional input vectors, and are applicable to efficient data compression. Our neural network contains a bottleneck layer with 500 nodes, while its input length depends on the number of MCs. Using this simple neural network, we were able to predict LFP waveforms from the *V*m dynamics of 1-5 MCs more accurately than expected by chance.

Few studies have recorded membrane potentials of MCs *in vivo* (Henze and Buzsáki, 2007; Soltesz et al., 1993). In particular, the *V*m dynamics of MCs during SWs have not been studied in detail. Only a single MC was recorded in our *in vivo* study. It is desirable to collect data from multiple MCs simultaneously for the same analysis as in *in vitro* experiments, but this is not technically feasible. However, the *in vivo V*m data may contain more information compared to the *in vitro* single MC data, because the nerve fibers are preserved without sectioning in the *in vivo* preparations. Indeed, the SW waveforms were predictable from the *V*m of MCs, which is consistent with the results of the analysis of *in vitro* samples.

Our analysis focused on SWs that could be predicted from individual MCs. On average, each MC accounts for about 10% of the SWs. This may be due to the fact that only a small number of MCs share the coding of SWs. We found that more than 30% of the SWs were predictable by at least five MCs. On the other hand, since MCs are biologically vulnerable (Scharfman, 2016), it is important that multiple MCs simultaneously take care of the same SW to avoid the risk of information loss due to neuronal cell loss. We attribute the sublinear rather than linear increase in the percentage of predictable SWs with increasing number of MCs to such overlapping MCs. We considered the sublinear increasing function as a general partial overlap problem and fit the data of **Figure 6C** to the exponential saturation function (*f* = 1 - α*^n^*) by the least squares method. As a result, we obtained the function *p* = 1 - 0.920 *^n^*, where *p* and *n* represent the ratio of predictable SWs to the total SWs and the number of MCs, respectively; the exponent α was 0.920 with a 99% confidence interval of [0.907, 0.933]. If the coverage of SWs by MCs assumes random sampling of SWs, the value of 1 minus α should be equal to the mean ratio of predictable SWs by a single MC, which can be obtained from **Figure 4A**. Indeed, the value was 0.080 (= 1 - 0.920), which was not statistically different from the mean ratio of predictable SWs (0.092 ± 0.048; *Z* = 0.250, *P* = 0.401, *Z*-test). Therefore, according to this approximation function, roughly 8 MCs can predict 50% of the total SWs, and 27 MCs can predict 90% of the SWs. Given that the hippocampal formation contains a total of 15,000 MCs, the number of MCs seems to be sufficient to cover the entire SW information (Amaral et al., 1990; Jinno and Kosaka, 2010). Therefore, the MC layer is likely to have both redundancy and independence in encoding information.

SWs originating in the CA3 region propagate to MCs and then to DG granule cells (Penttonen et al., 1997; Swaminathan et al., 2018); however, MCs are present in much smaller numbers than CA3 pyramidal cells and DG granule cells (Jinno and Kosaka, 2010). Thus, it is unclear how hippocampal information is efficiently encoded due to overcapacity in the MC layer. We observed distributed coding by MC populations. Furthermore, SWs with smaller amplitudes were processed by a smaller number of MCs, whereas SWs with larger amplitudes were covered by a larger number of MCs. In addition, we found that virtually all MCs responded reliably to each SW, suggesting that SW information is not sparsely encoded, but encoded by one or more MCs. Indeed, the use of more MCs correlated with better LFP prediction. This study not only provides important insights into the information processing of neural circuits at the bottleneck layer, but also introduces a new approach that employs machine learning to bridge different levels of time-series data, such as LFP and *V*m.

## Acknowledgments

This work was supported by JST ERATO (Y.I.: JPMJER1801), the Institute of AI and Beyond of the University of Tokyo (Y.I.), JSPS Grants-in-Aid for Scientific Research (Y.I.: 18H05525), Grant-in-Aid for JSPS Fellows (A.O.: 18J14574, 20J01255), and the JST CREST program (T.T.: JPMJCR23N2).

## Author contributions

A.O. and Y.I. designed the study; A.O. performed the experiments and data analysis; T.T. commented on analytical methods; A.O. and Y.I. wrote the manuscript.

## Methods

### Animal ethics

Animal experiments were approved by the Animal Experiment Ethics Committee of the University of Tokyo (approval number: P29-9) and performed in accordance with the University of Tokyo guidelines for the care and use of laboratory animals. These experimental protocols were carried out in accordance with the Fundamental Guidelines for Proper Conduct of Animal Experiment and Related Activities in Academic Research Institutions (Ministry of Education, Culture, Sports, Science and Technology, notice no. 71 of 2006), the Standards for Breeding and Housing of and Pain Alleviation for Experimental Animals (Ministry of the Environment, notice no. 88 of 2006) and the Guidelines on the Method of Animal Disposal (Prime Minister’s Office, notice no. 40 of 1995). All animals were housed under a 12-h dark–light cycle (light from 07:00 to 19:00) at 22 ± 1°C and had *ad libitum* access to food and water.

### Slice preparation

Acute slices were prepared from the medial to the ventral part of the hippocampus. On postnatal 21-29, male ICR mice were deeply anesthetized with isoflurane and decapitated. Their brains were rapidly removed and horizontally sliced (400 µm thick) at an angle of 12.7° to the fronto-occipital axis using a vibratome and an ice-cold oxygenated (95% O_2_, 5% CO_2_) cutting solution consisting of (in mM) 222.1 sucrose, 27 NaHCO_3_, 1.4 NaH_2_PO_4_, 2.5 KCl, 1 CaCl_2_, 7 MgSO_4_, and 0.5 ascorbic acid. This cutting angle preserved more Schaffer collaterals in the slices and was suitable for the reproducible generation of SWs (Mizunuma et al., 2014). Slices were incubated at 37°C for 1.0 h and maintained at room temperature for at least 30 min in a submerged chamber filled with oxygenated artificial cerebrospinal fluid (aCSF) containing (in mM) 127 NaCl, 3.5 KCl, 1.24 NaH_2_PO_4_, 1.3 MgSO_4_, 2.4 CaCl_2_, 26 NaHCO_3_, and 10 D-glucose.

### *In vitro* multiple patch-clamp recording

Recordings were performed in a submerged chamber perfused with oxygenated aCSF at a rate of 3-5 ml/min and a temperature of 33–35°C. Whole-cell current-clamp recordings were simultaneously obtained from up to five MCs in the dentate hilus, which was visually targeted using infrared-differential interference contrast microscopy. Patch pipettes (3-13 MΩ) were filled with a potassium-based solution consisting of (in mM) 135 potassium gluconate, 4 KCl, 10 HEPES, 10 creatine phosphate, 4 Mg-ATP, 0.3 Na_2_-GTP, 0.3 EGTA, 5 QX-314 and 0.2% biocytin. LFPs were recorded from the CA3c stratum pyramidale using borosilicate glass pipettes (0.5–1.2 MΩ) filled with aCSF. The signals were digitized at a sampling rate of 20 kHz. The data criterion for series resistance was less than 45 MΩ, and the change rate of series resistance before and after the recording was < 30%. The data were adopted when the mean resting potential was < −50 mV and when *Z-*scores of the mean membrane potentials for 30 s were between −2 and 2. MCs were verified by large whole-cell capacitances (> 45 pF) (Hedrick et al., 2017) and thorny excrescences on their proximal dendrites as determined by *post hoc* biocytin labeling (Murakawa and Kosaka, 2001).

### *In vivo* patch-clamp recording

Male ICR mice were used in the experiments on postnatal days 28-40. Before the surgery, the mice were exposed to an enriched environment for 30 min. The mice were anesthetized with urethane (1.95 g/kg, i.p.), and 1.0% lidocaine was subcutaneously applied to the surgical region. The anesthetized mice were fixed with a metal head-holding plate. The surgical procedures were described previously in detail(Matsumoto et al., 2016). Current-clamped recordings were obtained from MCs at depths of 940-1,150 μm from the dorsal alveus using borosilicate glass pipettes (3-8 MΩ). Patch pipettes were filled with a potassium-based solution consisting of (in mM) 135 potassium gluconate, 4 KCl, 10 HEPES, 10 creatine phosphate, 4 Mg-ATP, 0.3 Na_2_-GTP, 0.3 EGTA and 0.2% biocytin. Once a satisfactory whole-cell recording was obtained, the firing properties of the neuron were identified by applying a 500-ms current from −100 to +100 pA in increments of 20 pA. For each neuron, the spike responses to a brief inward current were examined. Cells were discarded when either the mean membrane potentials exceeded −40 mV or the firing patterns were fast-spiking. Tungsten electrodes (FHC, USA) were used for extracellular recordings from the hippocampal CA1 subregion. The LFP signals were amplified using DAM80 AC differential amplifiers (World Precision Instruments). All signals were sampled at 20 kHz using a Digidata 1440A (Molecular Devices). Tungsten electrodes were labeled with 1,1’-dioctadecyl-3,3,3’,3’-tetramethylindocarbocyanine (DiI, Invitrogen), and their tracks were visualized *post hoc*. Data were adopted when the electrode tips were placed within the CA1 region.

### Histology

For the visualization of patch-clamped neurons from the *in vitro* experiment, the slices were fixed in 4% paraformaldehyde and 0.05% glutaraldehyde at 4°C for more than 24 h. After the solution was washed out, the sections were incubated with 0.2% Triton X-100, a streptavidin-Alexa Fluor 594 conjugate (1:500) and 0.4% NeuroTrace 435/455 Blue Fluorescent Nissl stain (Thermo Fisher Scientific; N21479) overnight at 4°C. The tissue sections from the *in vivo* experiment were incubated with a rabbit primary antibody for GluR2/3 (Merck Millipore; AB1506; 1:200) overnight and then with a secondary goat anti-rabbit IgG (Cell Signaling Technology; #4412; 1:500) for 6 h. Fluorescent images were acquired using a confocal microscope (FV1200, Olympus or BX61WI, Olympus) and subsequently merged.

### SW detection

LFP traces were bandpass filtered at 2-30 Hz, and the SW peak times were determined at a threshold above the mean + 5×SD of the baseline noise(Mizunuma et al., 2014; Norimoto et al., 2018). The detected events were scrutinized by eye and manually rejected if they were erroneously detected. For the amplitude of an SW-triggered fluctuation in the subthreshold membrane potentials of MCs (Δ*V*ms), the mean *V*m (baseline) was determined 20 ms before the peak time of each SW and subtracted from the maximal *V*m within −30 to +40 ms from the SW peak time.

### Prediction of SW waveforms from *V*ms

A deep neural network model was designed to predict the SW waveforms from one to five simultaneously recorded *V*ms using Python 3.8. Our model has an encoder-decoder structure. The encoder compresses the input (*V*ms that correspond to SW) to a lower dimensional representation and extracts features from the input, whereas the decoder reconstructs the final output (SW waveforms) from the compressed vector. In the encoding operation, each *V*m trace for ±50 ms relative to the SW peak time (size 2001) was initially concatenated and passed through a fully connected layer. In this operation, the input was compressed to a feature matrix of reduced size (size 500). Our model was implemented using the Python deep learning library Keras and the TensorFlow backend. The network was optimized by adaptive moment estimation (Adam). The default values were used unless otherwise specified. Our model was trained to produce the original SW waveforms from *V*ms at the corresponding time points. To increase the number of datasets, we divided the quintuple, quadruple, and triple patch-clamp recording datasets into subsets with fewer MCs. Therefore, the analyzed data included predictions of the same SW waveform with different MC combinations. To assess the model performance on the entire dataset, 5-fold cross-validations were used. Each dataset was equally divided into 5 subsets; one subset was used as the test data, while the remaining 4 subsets were further divided into 10 subsets (one subset was used as the validation data, and the remaining 9 subsets were used as the training data). For each combination of inputs (*i.e.,* one *V*m, two *V*ms, three *V*ms, four *V*ms, and five *V*ms), the training lasted 100 epochs with a batch size of 16, and the RMSEs were calculated to assess how well the model predicted new data that were not used for training. As a randomized control, surrogate data were produced by shuffling the combinations of *V*ms; that is, the order labels of SWs were exchanged within each cell. For each dataset, 100 surrogates were produced, the model was also trained using these shuffled data, and the RMSE was calculated.

### Spatial entropy

For each spatial entropy predicted by a single MC, the prediction rate was computed from RMSE_real_ and RMSE_surrogate_, where RMSE_real_ and RMSE_surrogate_ represent the RMSEs between the original SW waveform and the SW waveforms predicted by deep neural networks trained using real *V*ms and surrogate *V*ms, respectively. SW was determined to be significantly predictive when RMSE_real_ were significantly lower than RMSE_surrogate_ calculated 100 times. For each SW and its four nearest neighbors in the MDS space, their five binarized prediction were averaged. The calculation was replaced by 1 if the SW was determined to be significantly predictive and 0.1 for otherwise. This procedure was repeated, and the averages were collected for all SWs. Then, the spatial bias of the mean prediction scores was quantified by spatial entropy, which was calculated as -Σ*P*i log_2_ *P*i, where *P*i is the probability frequency of the mean binarized prediction of MCs with the SW identity number *i*. The chance level of spatial entropy was estimated by 1,000,000 surrogates in which SWs were permeated among the MCs.

### Semicircular diagrams of the dentate gyrus

Data analyses were performed using ImageJ. The anatomical shape of the DG was normalized as a semicircular diagram to visualize the distribution of recorded MCs. The location of each MC was determined by its angle and distance in the DG as follows. For the definition of the angle of each MC, the edge of the CA3 pyramidal cell layer was defined as the center of the hilus, a line connecting the hilus center with the suprapyramidal blade (*i.e.*, the area closest to CA1) edge of the granule cell layer was defined as 0°, and the infrapyramidal edge was defined as 180°. For the distance of each MC in the hilus, the total distance from the edge of the CA3 pyramidal cell layer to the subgranular zone was determined, and the relative position of the MC was normalized to this distance. Using these methods, the angle and distance were determined for all MCs.

### Data acquisition and analysis

Data were analyzed offline using custom-made MATLAB routines (MathWorks, Natick, MA), and the summarized data are reported as the means ± standard deviations (SDs) unless otherwise noted. *P* < 0.05 was considered statistically significant.

### Data availability

All data generated or analyzed during this study are included in the manuscript. Original datasets and codes are provided on GitHub (https://github.com/ohchako/DistributedEncodingOfHippocampalInformationInMossyCells).

---

**Figure S1.**
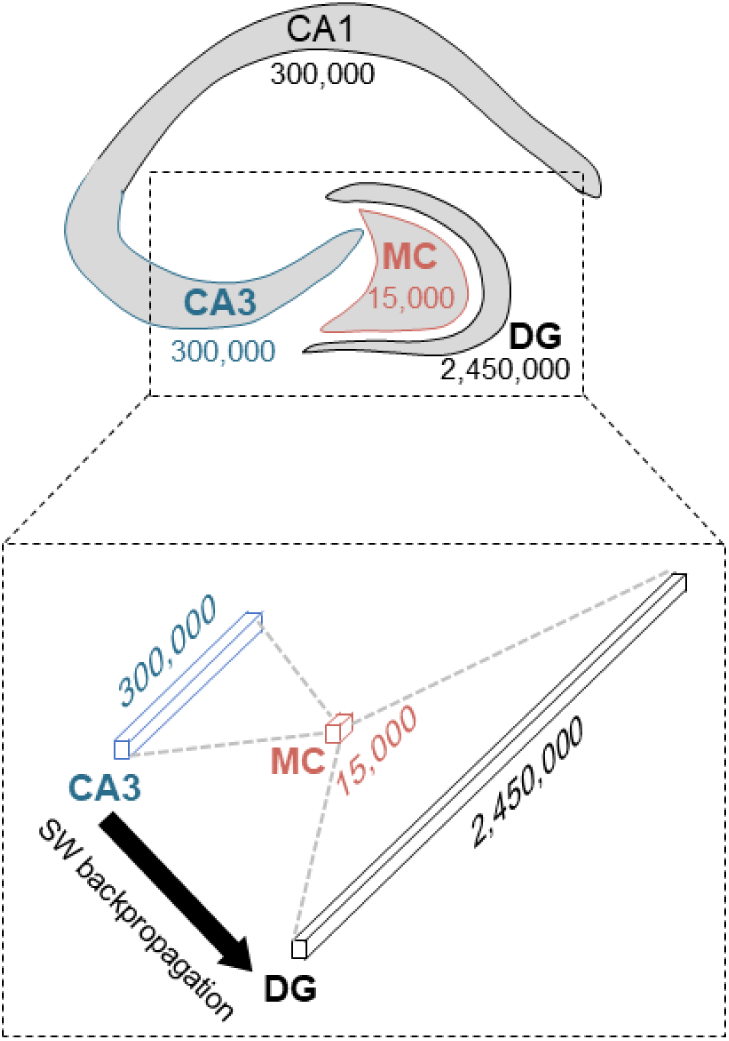
Estimated total number of excitatory neurons in each subregion in the rat hippocampus. Because the MC numbers were very low, the information had to be compressed in the MC layer during SW backpropagation from the CA3 region to the DG.

**Figure S2.**
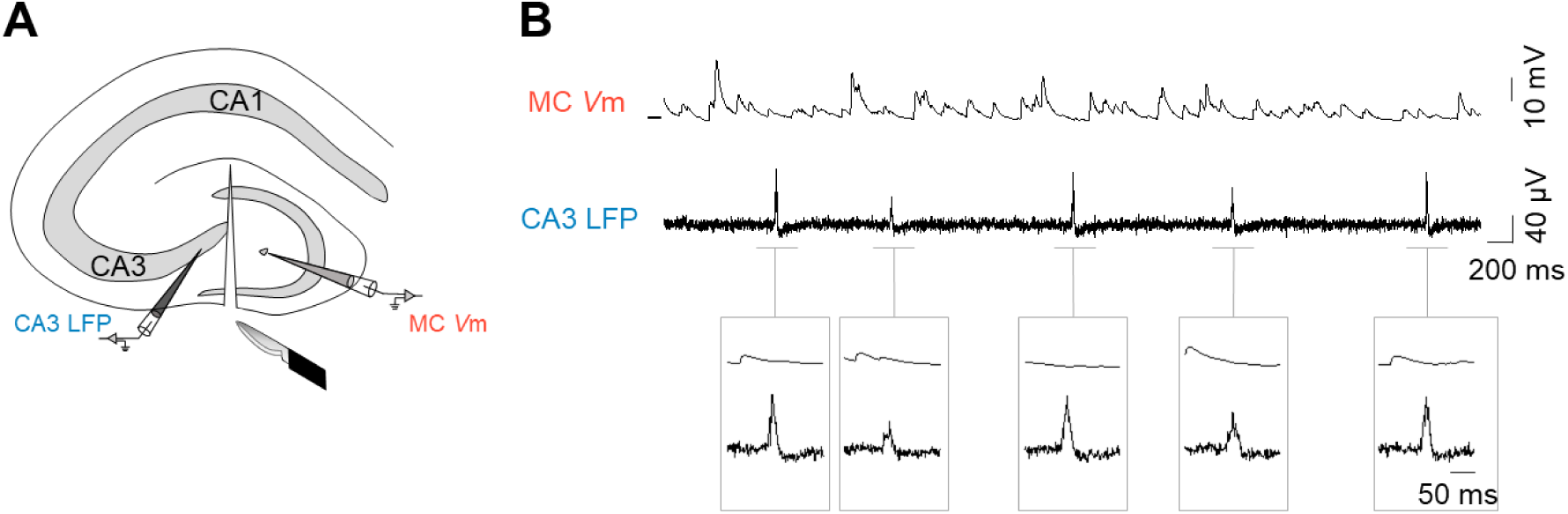
Lack of SW-induced depolarization of MCs isolated from the CA3 region. **(A)** The CA3 region and the hilus were surgically dissected before recordings. **(B)** A representative *V*m trace of a MC during SWs. The bar located on the left of the *V*m trace indicates the position of −60 mV.

**Figure S3.**
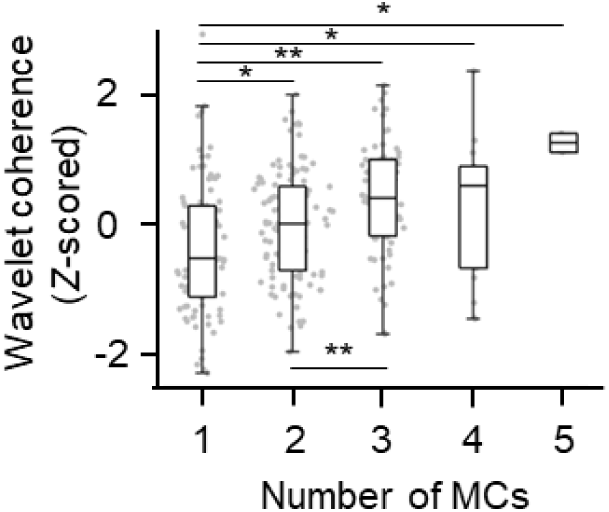
Reproducibility of high frequency (120-250 Hz) increases as the number of MCs increases. The wavelet coherence was computed between original SW traces and SW traces predicted from real *V*m datasets. The wavelet coherence values were averaged over the number of SWs and were *Z*-scored within each dataset. One cell: *n* = 28,463 SWs from 87 single MC recordings; 2 cells: *n* = 35,914 SWs from 113 MC pair recordings; 3 cells: *n* = 19,874 SWs from 67 trios; 4 cells: *n* = 4,403 SWs from 18 quartets; 5 cells; *n* = 271 SWs from 2 quintets, *F* = 7.84, *P* = 5.24 × 10^-6^, one-way ANOVA. \**P* < 0.05, \*\**P* < 0.005, Fisher’s LSD test.

**Figure S4.**
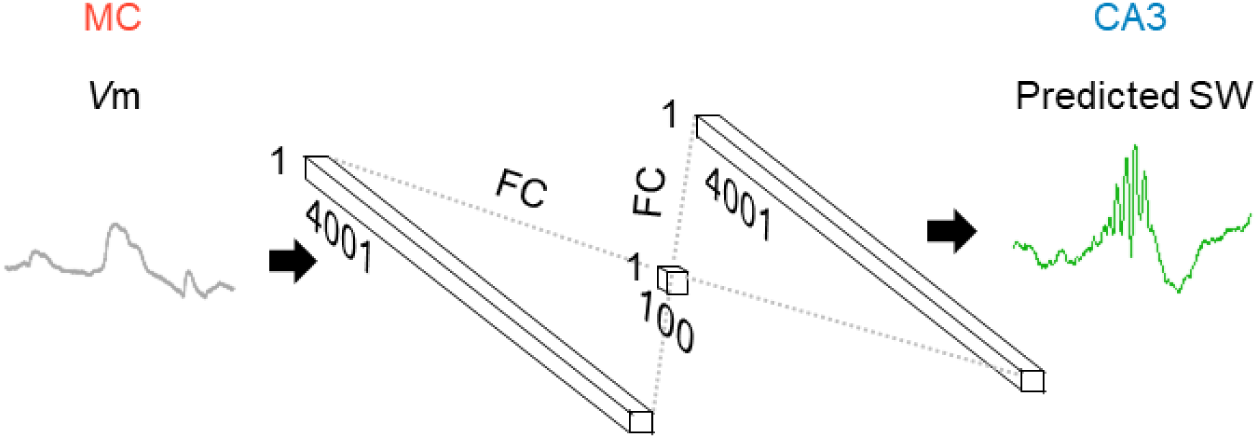
Design of *in vivo* neural network model. Input and output sizes were set to 4001 because the SW of *in vivo* has a longer SW than that of *in vitro*.

**Figure S5.**
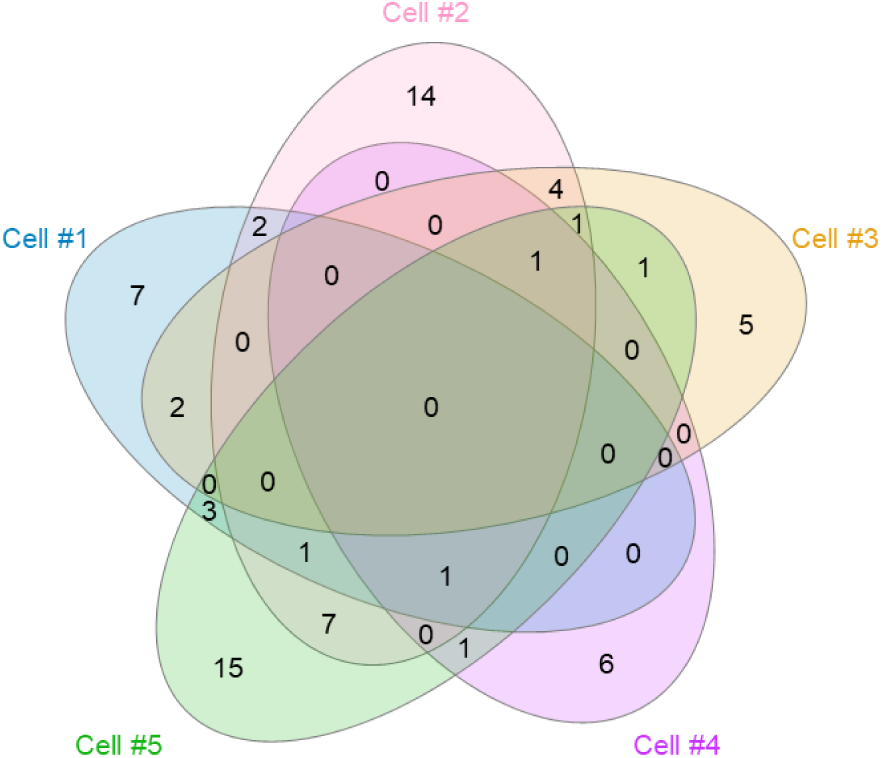
Visualization of overlapped SW. The number in each area represents the number of SW. In total, 178 SWs were recorded, with 71 predictable SWs, thus the total number of value is 71.

**Figure S6.**
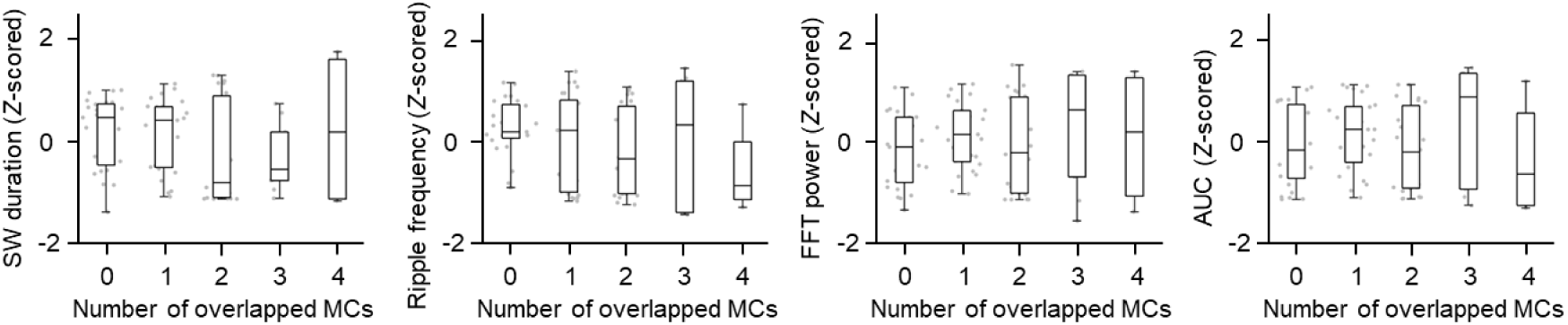
SW properties are maintained regardless of MC overlap. Correlations between SW properties and number of overlapped MCs. SW duration: *P* = 0.2249, *F* = 1.45; Ripple frequency: *P* = 0.2928, *F* = 1.26; FFT power (120-250 Hz): *P* = 0.3781, *F* = 1.07; AUC of FFT power (120-250 Hz): *P* = 0.4364, *F* = 0.96, one-way ANOVA. These values were obtained from 2 quintuple, 8 quadruple, and 15 triple recording datasets.

